# Animal acoustic communication maintains a universal optimum rhythm

**DOI:** 10.1101/2024.10.07.616955

**Authors:** T. Piette, C. Cathcart, C. Barbieri, K. M. Ming, D. Grandjean, B. Bickel, E.C Déaux, A-L. Giraud

**Affiliations:** Department of Basic Neurosciences, Faculty of Medicine, University of Geneva, Geneva, Switzerland; Department of Comparative Language Science, University of Zurich, Zurich, Switzerland; ISLE Center, University of Zurich, Zurich, Switzerland; Department of Evolutionary Biology and Environmental Studies, University of Zurich, Zurich, Switzerland; Department of Life and Environmental Sciences, University of Cagliari, Cagliari, Italy; Swiss Center for Affective Sciences, University of Geneva, Geneva, Switzerland; Université Paris Cité, Institut Pasteur, AP-HP, Inserm, Fondation Pour l’Audition, Institut de l’Audition, IHU reConnect, F-75012 Paris, France

## Abstract

Most animals interact with conspecifics through acoustic signals that are modulated in frequency and rhythm. While small animals vocalize at higher pitch than large ones due to the smaller size of their vocal apparatus, the rules governing vocalization rhythms throughout the animal kingdom remain unknown. Vocal rhythms serve as a natural information parser, and one possibility is that they are constrained by the neural rhythms of transmitter and receiver, known to be relatively conserved across species and independent of their size. In this study, we quantified acoustic rhythms across taxa and investigated their evolutionary history with regard to phylogeny and selective pressure. In 98 species from six classes, we tested the main factors likely to influence their communication rhythms: morphology, physiology, social complexity, mastication and detectability. Phylogenetic modeling did not confirm the influence of these species-specific factors, but rather point to a scenario where acoustic communication rhythms have been maintained around an optimum at around 3Hz in the biological (neuronal) delta range (1-4Hz) well before the mammals split. These results suggest that the rhythm of acoustic communication signals, unlike their pitch, has a universal neural determinant that has been conserved throughout evolution, allowing for intra- and cross-species signaling.

## INTRODUCTION

Acoustic signals allow for effective, instantaneous communication between individuals even at considerable distances. To be functional these signals’ structure must carry adaptive information with the importance of spectral features in doing so being well established^1–5^. Yet, acoustic signals are not only spectrally but also temporally structured, with rhythm having equally important communicative functions. This is particularly well exemplified by human speech, where speech rate is sufficient for comprehension^6,7^. In most languages, syllable production rate ranges between 4 to 9 syllables per second^8^. This frequency range corresponds to theta neural oscillations, which flexibly adapt to this rhythm during perception, such that modifying this flow impedes this process and decreases speech intelligibility. Rhythmic patterns allow the identification of syllables, words, and sentences and can help convey meaning, such as emphasis, intonation, and emotional state^9,10^. In animal calls, temporal features are no less important, for instance in vocal recognition^11^, mating behavior^12^, and predator avoidance^13^. Interestingly, calling rate is typically linked to arousal^1,14^ with high vocal rhythms eliciting rapid responses, possibly by targeting salience-related brain networks^15,16^.

Thus, the temporal patterning of acoustic sequences bears significant communicative function(s) and is not speech-specific. Yet questions remain as to what influences the evolution of rhythm across the animal kingdom. In this study we quantify rhythm across animal clades, test the most prominent hypotheses of signal structure evolution and build models of the most plausible evolutionary scenario. Four major selective forces may drive rhythm evolution. Firstly, the presence of theta rhythm in vocal productions and mouth movements of non-human primates suggests that this rhythm originates from the natural oscillatory movements of the articulators that are directly inherited from mastication^17–19^. Second, in animals that vocalize, morphological and physiological characteristics, such as breathing rate, heart rate, or metabolism could constrain rhythm range in an analogous manner to spectral features^20^. Thirdly, to be effective, acoustic signals must reach the receiver despite constraints from the living environment, which could thus influence not only spectral but also temporal features^21^. Additionally, the sociality-complexity hypothesis suggests a positive relationship between the complexity of the social environment and the complexity of a species’ vocal repertoire^22^. As rhythm would bear a direct relationship with the amount of information that can be transmitted in a given time unit, social complexity may also influence acoustic rate. Finally, it may be that none of these species-specific selective forces have influenced rhythm and that instead it has evolved through phylogenetic mechanisms of conservation and diversification that are either shared or diverse across lineages.

To test these different hypotheses, we quantified rhythm in acoustic sequences from birds, mammals, amphibians, insects, reptiles, and fish. Using Bayesian multilevel models for phylogenetic regression^23^, we controlled for phylogenetic relationships and evaluated, in accordance with the previously stated hypotheses, whether weight (as a proxy for breathing rate, heart rate, and metabolism), mastication status, sociality level or ecological characteristics could account for differences in rhythm. Finally, we compared phylogenetic models to assess the most plausible evolutionary scenario of rhythm across animals.

### Rhythm computation

We analyzed acoustic sequences from 98 species (58 birds, 28 mammals, 4 amphibians, 4 insects, 1 reptile, and 1 fish) to study the evolution of rhythm in animals. We calculated rhythm by analyzing variation in the signal amplitude, allowing broad applicability across species (Fig 1a, 1b, 1c, 1d). Validation against conventional methods showed consistent results (Fig 1e: F2.171=0.33, p=0.72) confirming the robustness of our method. Control of the impact of signal-to-noise ratio (SNR) and sequence length revealed no significant relationship with rhythm (Supp Fig 1a; Supp Fig 1b). Additional investigation of the allometric relationship between weight and dominant frequency (Supp Fig 2), and of the effect of context on rhythm (Supp fig 4a) further confirmed the validity of our vocal database.

**Figure 1.**
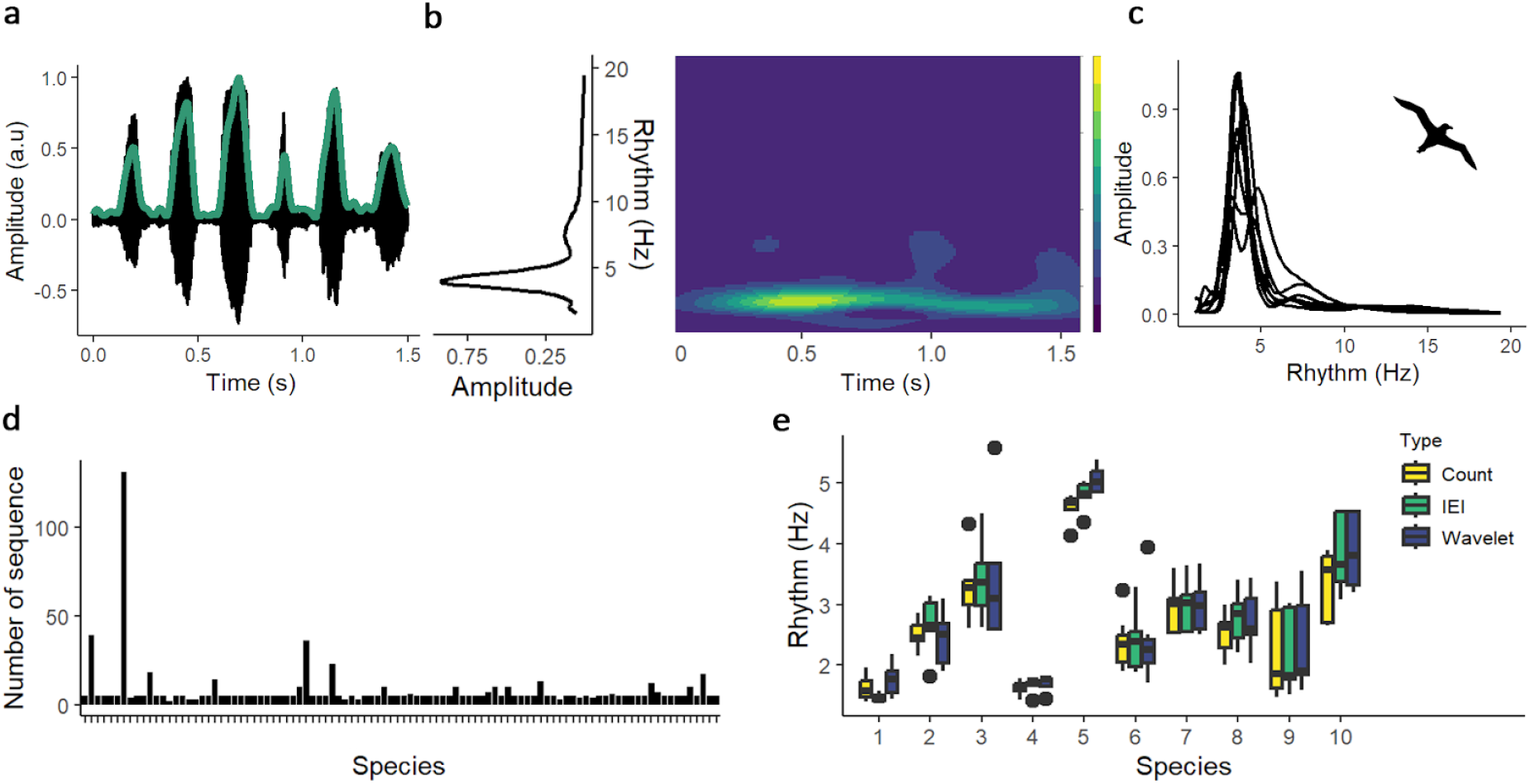
Methodology. a) Oscillogram of one acoustic sequence of polar skua call. b) Power spectrum and time frequency representation of the envelope of the previous sequence. c) Power spectra of the envelopes of all polar skua acoustic sequences b) Number of sequences per species (ordered by alphabetical order) c) Rhythm computation using sequences of 10 randomly selected species, employing, from left to right, number of elements per second (Count), inter-element intervals (IEI), and wavelet method (Wavelet).

### Species Specific Selective Pressure

To investigate which factors influence the evolution of rhythm, we fitted two phylogenetic regressions in a Bayesian multilevel framework A full model investigating the impact of weight, mastication status, and living environment while controlling for phylogenetic relatedness, and a null model controlling for phylogenetic relatedness only. We compared models via their leave-one-out expected log pointwise density (ELPD) and stacking weight. Due to heteroskedasticity, distributional (scale-location) models better fit the data, leveraging over 90% of the stacking weights (Sup Fig 3). Including the predictors did not improve predictive performance (Fig 2a) and the null model leveraged the highest stacking weight (Fig 2b). The posterior distribution of the regression coefficients of the full model revealed that none of our predictors had any decisive effect on rhythm, given that their 95% credible intervals (CI) all contain zero. We observed only very weak evidence for a negative effect of weight on rhythm in masticating species as zero was outside the 85% CI of their interaction coefficient (Fig 2c, 2d).

**Figure 2.**
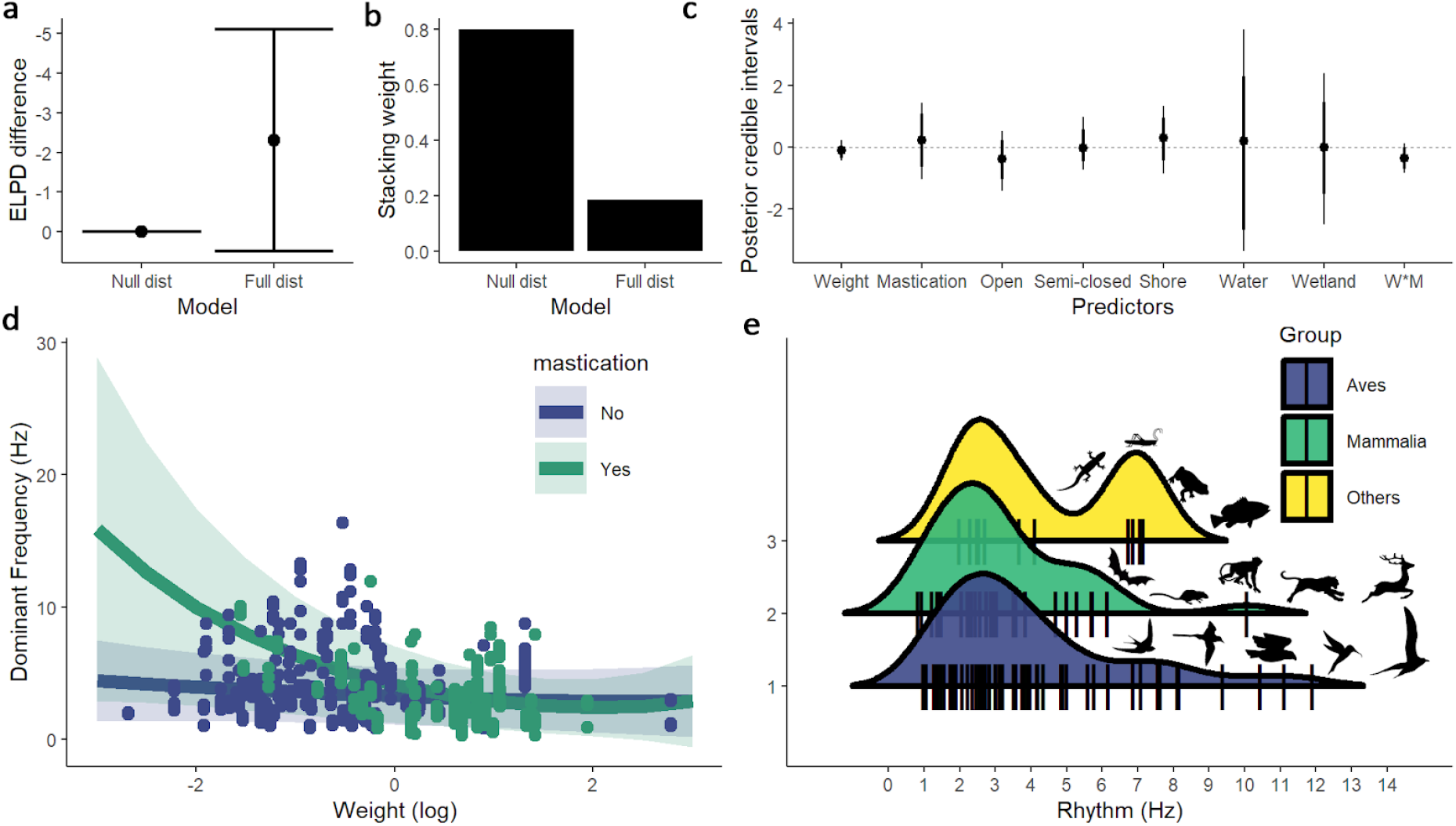
Bayesian multilevel model. a) Leave-one-out expected log pointwise density difference (ELPD) between the null and the full distributional (scale-location) models. b) Stacking weight of the models. c) Posterior credible intervals (95% and 85%) of the full distributional model (W*M = Weigh*Mastication) d) Rhythm plotted as a function of log-transformed weight with predicted slopes from the full distributional model and their standard error. e) Distribution of raw median rhythm in Hz in the different clades.

Taken together, these results indicate that inclusion of these predictors does not add decisive explanatory value to phylogenetic history. As vocal complexity, our indicator of social complexity, could not be included in the phylogenetic model due to unavailability for all species, we additionally regressed rhythm on vocal complexity (Sup Fig 4d: t=-0.75, p=0.46), revealing no relationship between the two.

### Phylogenetic History of Rhythm

Visual inspection of the median rhythm of our tested species shows that in every class, acoustic rhythm spans mainly the lower rates, in what is commonly called the delta range (Fig 2e). To understand if rhythm has randomly evolved in this range, favoring species-specific rhythm, or has been maintained around an optimum value, we fitted models of the evolution of vocal rhythm under Brownian Motion (BM) and Ornstein-Uhlenbeck (OU) processes ^24^, representing each evolutionary scenario respectively.

We fitted models of the evolution of vocal rhythm under BM and OU processes. Comparison of ELPD values and model stacking show that the OU model best fit the data (Fig 3a). The result is validated by models (Sup results) fitted separately for mammals and birds alone, with the OU model having the highest stacking weight (birds w_OU_ = 1, mammals w_OU_ = 0.925). The median posterior estimate of rhythm is 2.9Hz, with 95% CI of [.5, 5.1] Hz and 85% credible interval of [1.1Hz, 4.2] Hz (Fig 3b). This represents both the optimum to which the OU process reverts over time and the likely state at the root of the phylogeny. The posterior distribution of sigma represents the stochastic volatility (Fig 3c) and has a median of 0.69 and a 95% CI of (0.29 1.90). The posterior distribution of alpha represent the strength of attraction

**Figure 3.**
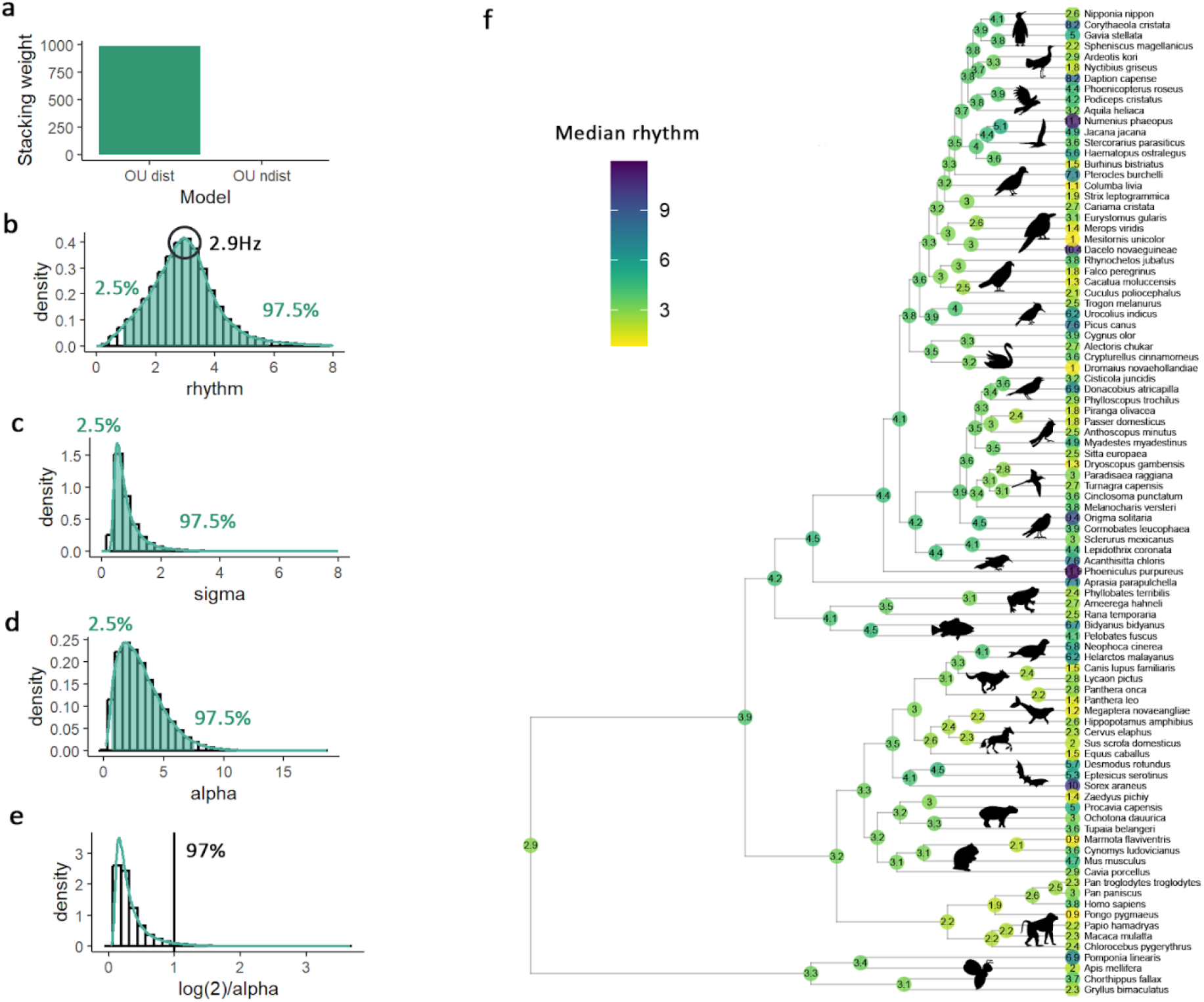
Phylogenetic modeling. a) Stacking weight of the OU and BM distributional models. b) Posterior distribution of rhythm in the OU distributional model. c) Posterior distribution of sigma, the scale of the drift process. d) Posterior distribution of alpha, the strength of selection. e) Posterior distribution of the phylogenetic half-life. f) Visualization of reconstructed median posterior values using a maximum clade credibility (MCC) tree.

(Fig 3d) and has a median of 2.79 and a 95% CI of (0.38 6.76). The proportion of posterior half-life estimates lower than 1 (height of the tree) is 97% (Fig 3e), supporting strong selection with fast reversals to the optimum. Consistent with this, rhythm values close to the optimum are also reconstructed for most interior nodes of the phylogeny (Fig 3f). In summary, the model suggests that there is a phylogeny-wide evolutionary pressure towards an optimal rhythm, to which species that deviate quickly revert to.

## DISCUSSION

Our finding of an animal-wide optimum rhythm for acoustic communication challenges the idea that it is shaped by biomechanical constraints. The latter would predict distinct rhythms between masticating and non-masticating species, along with a strong negative allometric relationship between rhythm and weight in masticating species. Instead, the weak effect of weight on acoustic rhythms observed in masticatory species indicates that biomechanical constraints only exert a marginal influence. By contrast, our analyses of dominant frequency (Supp fig 2) reconfirmed that spectral characteristics primarily depend on the specifics of the production organ, which chiefly vary with weight.^25^

With regards to the emergence of this optimum acoustic rhythm, its widespread evidence among birds and mammals suggests its presence in their last common ancestor, around 340 million years ago. Further, its existence in more remote species in our sample (insects, amphibians, fishes) speaks to even older roots (Fig. 3). Yet, the factors that contributed to the emergence of this rhythm and its persistence throughout evolution remain unclear. Given the diversity of hearing structures (cochlea, otic vesicle, tympanal organs etc.) and production modes (vocalization, stridulation, percussion etc.), it seems unlikely that this common rhythm reflects similarities in anatomical traits^26^. Likewise, given the diversity of living and social environments, such factors cannot be the main forces behind this conserved rhythm.

By contrast, basic neural mechanisms are remarkably conserved, and could therefore better explain a widespread optimum rhythm. On the production side, vertebrates share a common location for motoneurons responsible for vocalizations in the caudal hindbrain, believed to be involved in call duration and timing, supporting the claim of conserved mechanisms for acoustic rhythmic production^27^. In terms of auditory reception, this conserved rhythm around

2.9 Hz (85% interval 1.1Hz-4.2Hz) best matches delta brain oscillations (1–4 Hz), which have been observed across species, including mammals, reptiles, and insects^28–30^. Interestingly, delta oscillations are linked to active sensing, where organisms sample their environment to enhance perception/action cycles^31^. Slow oscillations help integrate sensory information and are particularly suited for integrating slow-varying acoustic cues, that are usually the signature of the emitter’s identity, e.g. vocalic cues^32^. In sum, this production rhythm best matches a neural rhythm that is important for assigning significance to acoustic signals, which would thus result in an effective communicative design.

Yet, speech perception research has revealed that auditory processing relies on (at least) a two-timescale processing^10^. Similarly, our control analyses on context show that different call types operate within this conserved rhythm (Sup Fig 4), requiring a faster analysis window to be resolved. As such, it is possible to envision a dual receptive strategy relying on slow analyses for signal identification, and fast analyses for signal detectability and discrimination, useful for e.g. distinguishing different call types^33–36^. Though this report does not demonstrate the fast timescale’s evolution, an interesting avenue would be to probe for an optimum that combines fast and slow scales during acoustic communication.

Finally, these results suggest another interesting aspect. The maintenance throughout evolution of a slow rhythm for acoustic signal production and perception de facto results in a common communication channel across coexisting species. The common low-range (delta scale) could thus permit cross-species identification and offer possibilities for interspecies signaling (e.g. a common danger) and/or eavesdropping, conferring evolutionary advantages.

In summary, spectral and temporal features, both key to effective communication, evolve following different trajectories. While spectral traits diversify along with the hearing organ anatomy, rhythm may be primarily shaped by basic neural factors, responsible for both production and perception processes and which lead to a conserved optimal rhythm.

## METHOD

### Vocal sequences

To perform an extensive phylogenetic comparison on a balanced phylogeny, we collected acoustic and biological data for at least one species per infra-order of tetrapods, when data were available, as well as a few species of insects and fishes, to obtain a good representation of rhythm throughout the phylogeny. Acoustic sequences were gathered from public and private databases (Fonozoo, Cisro, Berlin Museum fur Naturkunde and Xeno-canto), online videos platform (Youtube, Dailymotion), and from different research groups that kindly shared audio files.

### Weight

Weight presents an allometric relationship with morphological features and physiological processes involved in acoustic productions across various species. Thus we collected biological data on the mean weight for each species, to serve as a proxy for heart rate, breathing rate and metabolism. This involved calculating the average weight by considering both the minimum and maximum weights recorded for each species, irrespective of sex. These data were primarily obtained from the handbook of mammals of the world^37^, and the handbook of birds of the world^38^. When data were unavailable from these sources, we looked for reference articles.

### Beak Size

Just as masticatory abilities may have influenced rhythm in mammals, a similar proposition could be made regarding beak morphology in birds. Unlike other animals, the morphological traits of a bird’s beak do not consistently adhere to an allometric relationship with its weight^39^. Thus we collected information on beak length, width and depth of our species. As these measures were not available for all species, we could not include them in the phylogenetic model. We therefore built an additional linear mixed model investigating the variation of rhythm, including group and order as random effects, and beak length, width, depth, and their interactions as fixed effects. Data are available in the study github.

### Living environment

As environmental conditions can impact vocal communication^21^, we also collected data on the typical habitat of each species. We used a five level categorical classification, with habitats being either classified as closed (defined as habitats with heavy tree coverage), semi-closed (defined as habitats with light three coverage or human cities), open (defined as fully open habitat with no three coverage or obstacle), shore (for species living near a significant amount of water such as lakes, rivers or seas) or water (for species living below the water surface). These data were primarily obtained from the handbook of mammals of the world, and the handbook of birds of the world. When data were unavailable from these sources, we looked for reference articles. Data and linked references are available in the study github.

### Mastication status

As some have proposed that rhythmic communication in vocalizing animals may be linked to mastication regime^17,19^, we also classified each species according to their mastication status (yes or no).

### Social complexity

As social complexity increases, individuals may need to communicate more information in a given time, and therefore speed up their communication. As communication signals have been linked to social complexity^22^, we also gathered species vocal repertoire complexity (number of distinct calls in the species vocal repertoire) when this information was available. As this measure was not available for all species, we could not include it in the phylogenetic model. We therefore built an additional linear mixed model investigating the variation of rhythm, including group and order as random effects, and vocal complexity as fixed effect. These data were primarily obtained from the handbook of birds of the world, and reference articles. Data and linked references are available in the study github.

### Acoustic sequence selection and pre-processing

Based on a cross-species literature search^40–43^, we defined sequences as recordings of acoustic displays emitted by a single individual, containing more than two calls separated by less than two seconds of silence. For computational purposes, we selected only recordings lasting more than one second. We included in our analysis species for which we had at least five different sequences from five different individuals. Species with fewer sequences were included if they were the only available representatives of their infra-order, resulting in a database of 98 species, including 58 birds, 28 mammals, 4 amphibians, 4 insects, 1 reptile, and 1 fish.

### Rhythm analyses

To quantify rhythm in these acoustic sequences, we decided to adapt the method developed by Tilsen et al to compute rhythm in human speech production^44^. This method uses the signal amplitude to automatically compute the rhythmic component of a sequence, without making any assumption on the components’ size, and is thus widely applicable across all species regardless of variations in unit size or spectral characteristics. First, we denoised the sequences using a first-order Butterworth filter, with a bandpass filter between a minimal frequency (minF), defined as 200 Hz below the minimum frequency of the animal call, and a maximum frequency (maxF), defined as 200 Hz above the maximum frequency of the animal acoustic signal obtained from reference articles. When this information was not available, we applied a large range filter with a 100 Hz minF and 10000 Hz maxF. We then computed the normalized envelope of the denoised sequences using the Hilbert Transform. Next, we low pass-filtered this envelope with a fourth-order Butterworth filter with a 20 Hz cut-off frequency to obtain the slow changes in acoustic energy. Before further analysis, we downsampled the resulting signal at 150 Hz for computational purposes, and then applied a continuous wavelet transform using the Morlet wavelet to obtain a time-frequency representation of the amplitude envelope. We replaced the Fourier transform with a wavelet transform, to allow for more flexibility with regards to the variation in sequence length present in our dataset. We finally analyzed that representation’s power spectrum to extract the five frequency peaks of highest amplitude in the power spectrum and used the time-frequency representation to select the main rhythmic component conserved across the entire sequence.

To assess the validity of the proposed methodology, we conducted a comparative analysis between the calculated vocal rates of a subset comprising 10% of our database and those derived from two widely accepted conventional approaches: 1) by counting the number of elements per second (Count) and 2) by computing the inter-element interval (IEI). The three methods gave sensibly similar results (Figure 1d: F_2.171_=0.33, p=0.72), hence validating our rhythm quantification method.

### Signal to noise ratio (SNR) and Length

As further control analyses, we quantified recording durations in seconds and signal-to-noise ratio in decibels. To control for the effect of both factors and their interaction on rhythm, we build a linear mixed model investigating the variation of rhythm including group and order as random effects, and SNR, length and their interaction as fixed effects.

### Dominant frequency analyses

To determine dominant frequencies, which unlike the fundamental frequency are measurable in all types of communicative signals^25^, we isolated the first acoustic unit in each denoised sequence. We then applied a single discrete Fourier transform to compute the power spectrum of these units, and extract the peak of highest amplitude. The obtained results were also visually controlled in Praat, to make sure that the extracted dominant frequency matched the acoustic energy present in the unit. If the first unit had poor signal-to-noise ratio leading to inaccurate computation of the dominant frequency, we selected the next unit in the sequence.

### Context effect control

While the existing literature highlights the importance of context and its correlated arousal levels on acoustic signal rate^1^, for most species we were not able to obtain these data. Nevertheless, whenever possible we selected recordings of different call types for each species. Further, we performed separated analysis of variance (ANOVA) to control for the effect of call type on rhythm in three species: one avian and two mammalian, from which we could obtain different call types, including contact calls, alarm calls, songs, agonistic and antagonistic vocal displays.

### Phylogenetic tree sample reconstructed from genetic sequences

To test our hypotheses of interest we used phylogenetic comparative methods (PCM), a broad family of methodological tools for characterizing and controlling for the evolutionary dynamics thought to give rise to the data under study. PCMs require a representation of the relatedness of the taxa under study in the form of a phylogenetic tree sample. To represent the tree topology of the species we performed a phylogenetic analysis based on comparable genetic sequences, using a Bayesian framework to infer a posterior tree sample. We first matched each species in the sample with their closest genetic proxies in GenBank^45^, and extracted mitochondrial DNA for the corresponding species. For 54 of the 98 species, matching mitochondrial genomes were available from literature and deposited in GenBank. For the 44 remaining species we chose proxies from another closely related species. We took a species within the same genus when possible; if this was not available, we chose species within the family of the target species, after confirming that no more than one species per family was included in the original list. Only for four target species - *Correlophus ciliates* (Squamata), *Galbula ruficauda* (Aves), *Leptosomus discolor* (Aves), *Phaethon rubricauda* (Aves) - we did not find proxies within the family and we had to find a proxy within the order. To choose the best mtDNA proxy with those deposited in GenBank, we considered completeness of the available mitochondrial sequences and comparable average size, weight and environment of the target species. Maximum missing data is 100 base pairs, for an average size of 16706 base pairs. MtDNA genomes were aligned with MAFFT software^46^ and standard settings. The alignment was manually screened in BioEdit (version 7.2, https://thalljiscience.github.io/) for spotting irregularities and potential outlier sequences. Sequences were then cut to keep only the coding region, which is more conserved across species, using the *Homo sapiens* sequence as a reference. The final alignment consisted of 21860 base pairs, which include large INDELs sections to accommodate alignment between the most divergent species (e.g. *Apis mellifera*).

We used BEAST2 to generate the trees, running 10’000’000 iterations of Markov chain Monte Carlo (MCMC) with a thinning interval of 1000. We used the following settings to approximate the broad evolutionary range of the species considered: assuming an HKY substitution model, a strict clock (Uniform rates across branches), and a Birth-Death tree prior with a Yule birth rate. This resulted in 10’000 trees, of which we use 50 for phylogenetic comparative analyses.

### Bayesian multilevel models for phylogenetic regression

To assess the impact of several predictors of interest on different properties of variation in (1) vocal rhythm and (2) dominant frequency, we used phylogenetic regression modeling, a comparative method that assesses the effect of predictors on a response while controlling for the phylogenetic relatedness of the taxa. Due to heterogeneity in the number of datapoints and individuals in each species, we employed both non-distributional regression models, which model the mean of the response variable as a function of predictors, and distributional (scale-location) regression models, which model both the mean and standard deviation of the response variable as a function of predictors. We control for species-level idiosyncrasies in both the median and (in some cases) standard deviation of rhythm via phylogenetic random intercepts and slopes (phylogenetic random effects are similar to the standard random effects used in hierarchical regression modeling, but are generated by a Gaussian Process with a covariance kernel that is a function of the phylogenetic patristic distances between species under study rather than independently and identically distributed with diagonal variance).

We fitted four phylogenetic regression models using brms^23^ for each response variable (vocal rhythm, dominant frequency), resulting in eight models. These predictors consisted of average species weight in kilograms, which was log-transformed, centered around zero, and standardized^47^, mastication status, and living environment of the species. We also consider the interaction between mastication status and weight.

Our distributional models have the following basic generative process (below, α^μ^+β*X*_*i*_ is shorthand for all model predictors, including fixed and random effects):

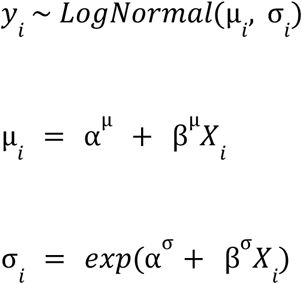

Non-distributional models have the following structure:

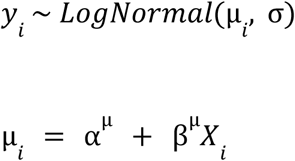

We employ the default priors of brms.

The first of the four models was a full distributional one that modeled both the expected median and variance of the response variable as a function of these predictors, while controlling for species-level idiosyncrasies in both the median and variance of rhythm via phylogenetic random intercepts and slopes. The second was a full non-distributional model that treated only the expected median rhythm as a function of the predictor variables as well as phylogenetic random intercepts and slopes. The third of these was a null distributional model that included only phylogenetic random intercepts and slopes for the expected median and variance. The final model was a null non-distributional that included only phylogenetic random intercepts and slopes for mean rhythm. We ran each of these models for 4000 iterations of the no U-turn sampler over 4 chains with a log-normal link function and discarded the first half of samples, aggregating posterior samples across the retained sampled trees.

Models in brms have the following formulae:

Full, distributional

~~~
bf(frequency ∼ weight*mastication + environment + (1 +
weight*mastication + environment | gr(taxon, cov =
phylo.cov))
~~~

~~~
sigma ∼ weight*mastication + environment + (1 +
weight*mastication + environment | gr(taxon, cov =
phylo.cov)))
~~~

Full, non-distributional

~~~
bf(frequency ∼ weight*mastication + environment + (1 +
weight*mastication + environment | gr(taxon, cov =
phylo.cov)))
~~~

Null, distributional

~~~
bf(frequency ∼ (1 | gr(taxon, cov = phylo.cov)), sigma ∼
(1 | gr(taxon, cov = phylo.cov)))
~~~

Null non-distributional

~~~
bf(frequency ∼ (1 | gr(taxon, cov = phylo.cov))
~~~

We compared fitted models via their leave-one-out expected log pointwise density (ELPD) values^48^ and stacking^49^, which average predictive distributions of different models to generate weights representing their relative predictive power. We used the function loo_compare to measure differences in ELPD across models. Finally, We inspected posterior distributions of regression coefficients of the full distributional model to assess the effects of predictors of interest.

### Evolutionary dynamics of vocal rhythm

We further investigate the properties of rhythm across species using two Gaussian Process models of continuous trait evolution, asking specifically whether the evolution of vocal rhythm is characterized by a random process of drift (characterized by Brownian motion) or whether selective forces draw rhythm values toward an optimal value over time (a mean-reverting scenario characterized by an Ornstein-Uhlenbeck process). Under Brownian motion, the displacement of a continuous trait at time *s* has a variance proportional to the amount of time elapsed over the course of displacement (below denoted as *t*), where σrepresents the scale of the drift process:

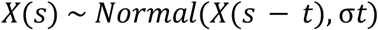

Under an OU process, the displacement of a character has the following formula:

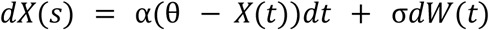

In the first component of the sum, α represents the strength of selection to the optimal value θ. The second component represents a process of Brownian motion, with σ representing the scale of drift. Thus, the OU process allows for both selective and random forces in character evolution.

An standard way to interpret α is to transform it to the phylogenetic half-life, ln 2/α^50^. This is interpreted as the average time for a trait to evolve halfway from an ancestral state toward a new optimum, indicating how long it will take before adaptation to a new regime is more influential than constraints from the ancestral state. If half-life values are greater than the height of the phylogeny (1 in our case, as the tree length is scaled to unit height), the process increasingly resembles Brownian motion and involves a slower adaptation speed.

As above, we employ a distributional approach, allowing species-level mean rhythm values and species-level standard deviations of rhythm values to evolve over the phylogeny according to BM or OU processes.

The distributional BM process has the following generative process:

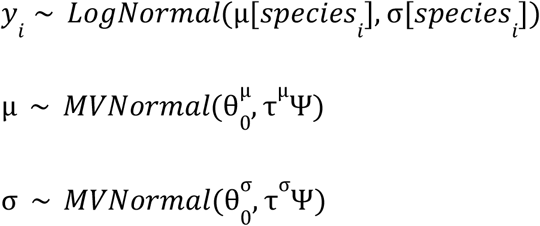

θ_0_ represents the trait value at the root of the tree, while Ψ is a matrix of the shared history (the time between the root age of the tree and the most recent common ancestor) of each pair of nodes in the tree and τ is the positive scale of drift. Conventions are as above.

The distributional OU process has the following generative process:

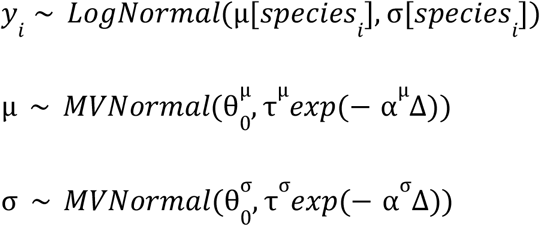

θ_0_ represents the trait value at the root of the tree, τ is the positive scale of drift, *α* is the positive strength of selection, and Δ is a matrix of pairwise cophenetic distances between species in the phylogeny, scaled to a maximum distance of 1. Conventions are as above. We place *Normal(0,1)* priors over unconstrained parameters and *Gamma*(1,1) priors over positive parameters. In addition to running these models on all species in our sample, we validate results by running models on bird and mammal species alone.

### Phylogenetic reconstruction of rhythm values

Rhythm values were reconstructed to internal nodes of the maximum clade credibility (MCC) tree of the phylogeny, using ggtree package^51^ by drawing ten draws from each of the posterior distributions inferred from the 50 different trees in the tree sample and sampling values at internal nodes of the tree from the normal distribution parameterized by the OU process, conditioned at the expected tip values (fig 3f).

### Software

All analyses and visualization were done using Stan and R version 4.1.2 (2021-11-01) with the following packages Seewave^52^, Soundgen^53^, DoBy^54^, Lme4^55^, MuMYn^56^, brms^23^, ggplot2^57^.

## Supporting information

Supplementary results

## Data and code availability

All data and code used for the analysis in this study are available in a public GitHub repository. This repository contains the raw data, scripts for data processing, analysis, and visualization, as well as detailed instructions for reproducing the results. The repository can be accessed at https://github.com/chundrac/phylo-acoustic-rhythm.

## ACKNOWLEDGMENTS

The NCCR Evolving Language, Swiss National Science Foundation Agreement Nr. #51NF40_180888, funded this work. The authors are grateful to the CSIRO Australian National Wildlife Collection, (https://ror.org/059mabc80), Fonoteca Zoologica (https://www.fonozoo.com/), the Museum für Naturkunde of Berlin (https://www.museumfuernaturkunde.berlin) and the Xeno-Canto Foundation for Nature Sounds (https://xeno-canto.org) for their assistance in undertaking this research. The authors thank all the researchers from the NCCR Evolving Language as well as Pascal Belin and Steffen R.Hage for their insights on the project. The authors would like to thank Aaron Bauer, Andrew Spencer, Alex Kwet, Alex Rohtla, Camila Ferrara, Daniel Blumstein, Élodie Briefer, Émilie Genty, Frank Lambert, Isabelle Charrier, Gerald Carter, Hans Schneider, Marc D. Hauser, Nikola Falk, Peter Boesman and Robert Seyfarth for providing us with recordings for this study. ALG is funded by the Fondation pour l’Audition (GA-FPA IDA11).

## CONTRIBUTIONS

T.P, E.D, D.G, and A-L.G conceptualized the project. T.P conducted the acoustic data collection, while K.K.M was responsible for the genetic data collection. The acoustic analysis methodology was developed by T.P and E.D, with T.P conducting the acoustic data analysis.

C.B constructed the phylogenetic tree. Phylogenetic analyses were designed by C.C and B.B and carried out by C.C. T.P wrote the initial draft, with C.C providing detailed methodological input on the phylogenetic analysis. All authors contributed to the review and editing of the manuscript. E.D and A-L.G provided supervision. A-L.G provided resources, and funding for the project.

