## Supplementary results for "Animal acoustic communication maintains a universal optimum rhythm"

**Affiliations:**

### SUPPLEMENTARY RESULTS

### Noise and Length Control :

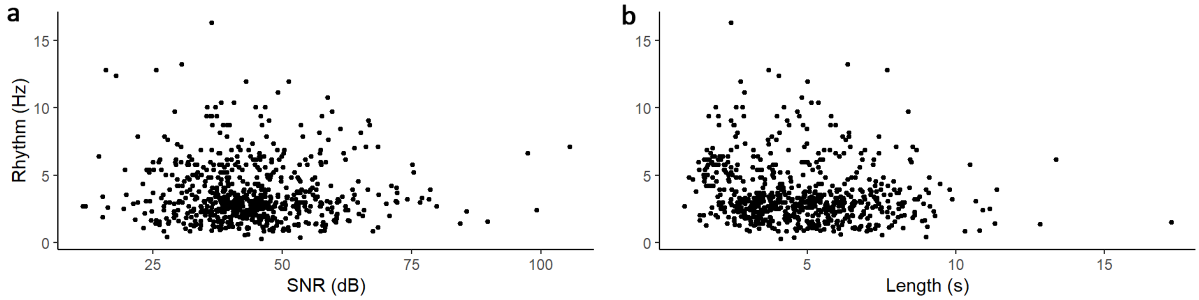


**Supplementary Figure 1 : Signal to Noise Ratio and sequence length effect on rhythm** a) Rhythm (Hz) as a function of signal-to-noise ratio (SNR) showing the absence of relationship between SNR and Rhythm ($t=-0.21$, $p=0.84$, $R^2=0.02$) b) Rhythm (Hz) as a function of recording length showing the absence of relationship between sequence length and rhythm ($t=-1.45$, $p=0.15$, $R^2=0.02$).

#### Phylogenetic regression of Dominant Frequency (DF)

As our acoustic data were gathered from both supervised and unsupervised dataset, we decided, as a validation procedure, to check for the presence of the well established allometric relationship between weight and dominant frequency in our dataset. To do so, we fitted two phylogenetic regressions using Bayesian multilevel models. A full model investigating the impact of weight, mastication status, and living environment while controlling for phylogenetic relatedness, and a null model controlling for phylogenetic relatedness only. Due to heterogeneity in the number of datapoints and individuals in each species, we employed both non-distributional and distributional regression models. The full distributional model with predictors included has the highest ELPD value (Sup Figure 2a). In terms of stacking weight, we see that the full distributional models with predictors has the majority of the stacking weight, but the null distributional model still has a substantial amount of stacking weight (Sup Figure 2b). We inspect the credible intervals (CIs) of posterior distributions of model coefficients of the the full distributional model to determine which predictors have an effect on dominant frequency, the response value. 95% and 85% CIs for model coefficients are found below, along with the proportion of posterior samples for which the coefficient value is greater or less than zero. We take coefficients whose 95% CIs exclude zero to represent decisive evidence for an effect, and coefficients whose 85% CIs exclude zero to represent strong evidence for an effect. As is clear, there is a decisive negative effect of weight on dominant frequency, and strong evidence for a negative interactive effect between mastication and weight on dominant frequency (Sup Figure 2c, 2d), thus validating the accuracy of our database.

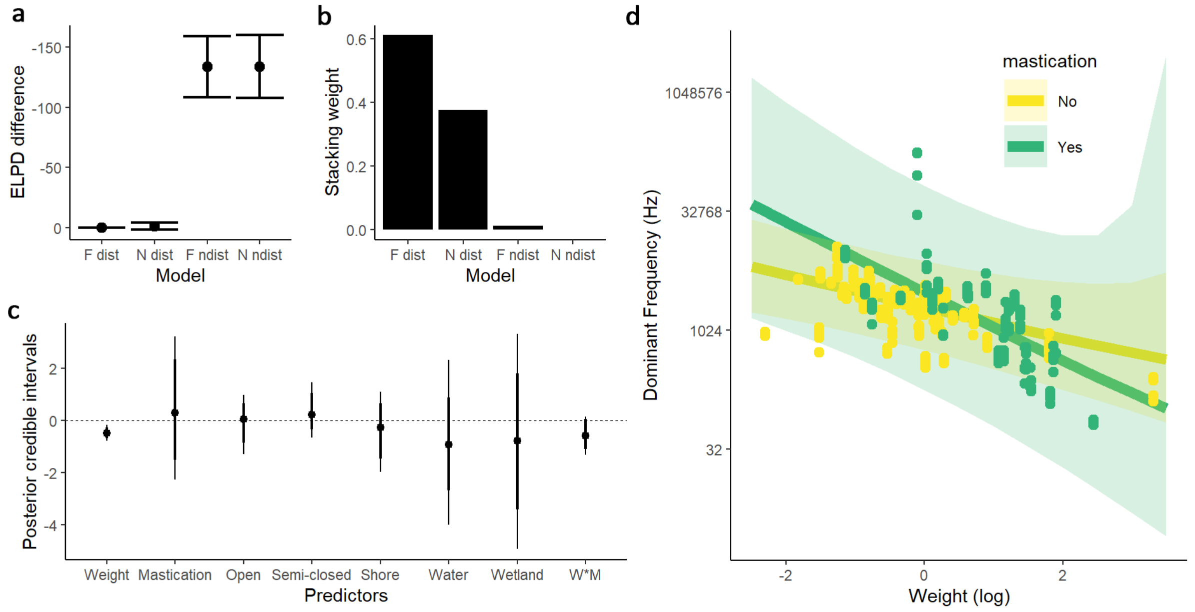

**Supplementary Figure 2. Dominant Frequency Phylogenetic Regression** a) Leave-one-out expected log pointwise density difference (ELPD) between the full and the null distributional models. b) Stacking weight of the models. c) Posterior credible interval (95% and 85%) of the full model d) Dominant frequency plotted on a logarithmic scale as a function of log-transformed weight with predicted slopes from the full distributional model and their standard error.

**Phylogenetic regression of Rhythm :**

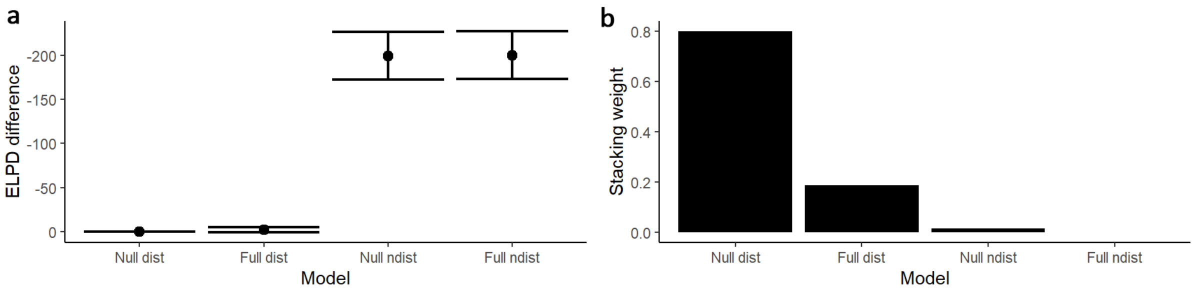


**Supplementary Figure 3. Rhythm Phylogenetic Regression** a) Leave-one-out expected log pointwise density difference (ELPD) between the null distributional ("dist") model and the others ("ndist" for "non-distributional, modeling only the mean") . b) Stacking weight of the models.

**Phylogenetic History of Rhythm in birds and mammals**

Independent analyses of the evolution of rhythm in birds and mammals confirm the results on the entire dataset, with the OU model having the highest stacking weight (birds = 1, mammals = 0.925). This indicates that evolutionary history of rhythm is consistent across clades, showing conservation of rhythm and its constraints across the different groups.

### Context

As the existing literature highlights the importance of context and its correlated arousal levels on acoustic signal rate, we investigated the effect of context on acoustic rhythm in three distinct species. Analysis of variance (ANOVA) of the effect of call type on rhythm in baboons (*Papio anubis*) revealed significant effect of call type on rhythm ($F_{2,27} = 14.99$, p=0.0004) with post-hoc tukey tests showing significant differences in rhythm between scream and affiliative grunt, as well as between threat grunt and affiliative grunt (Sup fig 4a). Similar analysis in European stone-curlews (*Burhinus oedicnemus)* and dogs *(Canis lupus familiaris)* showed comparable results ($F_{2,21}$=4.546, p=0.02; $F_{4,136}$=3.632, p=0.007), with call type having a significant impact on rhythm. However, the median rhythm for each call types are contained between ± 1Hz around the median rhythm of the species (Sup fig 4a,4b,4c). This illustrates that, even though context bears a significant influence on vocal rate, this influence is limited to a narrow and constrained range around the median vocal rate of the species.

### Vocal complexity

As vocal complexity is linked to social complexity [7], we build a linear regression model between rhythm and vocal complexity, to investigate the effect of social complexity on rhythm in animals. Vocal complexity did not show any significant relationship with rhythm (t=-0.748, p=0.46). The model failed to account for a significant proportion of variance in rhythmic behavior ($R^2$=-0.01, p=0.46), suggesting that other factors beyond social complexity may influence rhythmic behavior (Sup Figure 4d).

### Beak size

To investigate the effect of beak morphology on rhythm in birds, we build a linear regression model examining the relationship between rhythm, beak length, width, depth, and their interactions. Beak length (t=-1.4, p= 0.17), beak width (t=-0.98, p= 0.33), and beak depth (t=0.35, p=0.73) did not show any significant relationship with rhythm (Sup Figure 4e,4f,4g). Similarly, interactions between these variables were not significant predictors of rhythmic behavior (t=-0.41, p=0.68). The model failed to account for a significant proportion of variance in rhythmic behavior ($R^2$=-0.01, p=0.5), suggesting that other factors beyond morphological traits may influence bird rhythmic behavior.

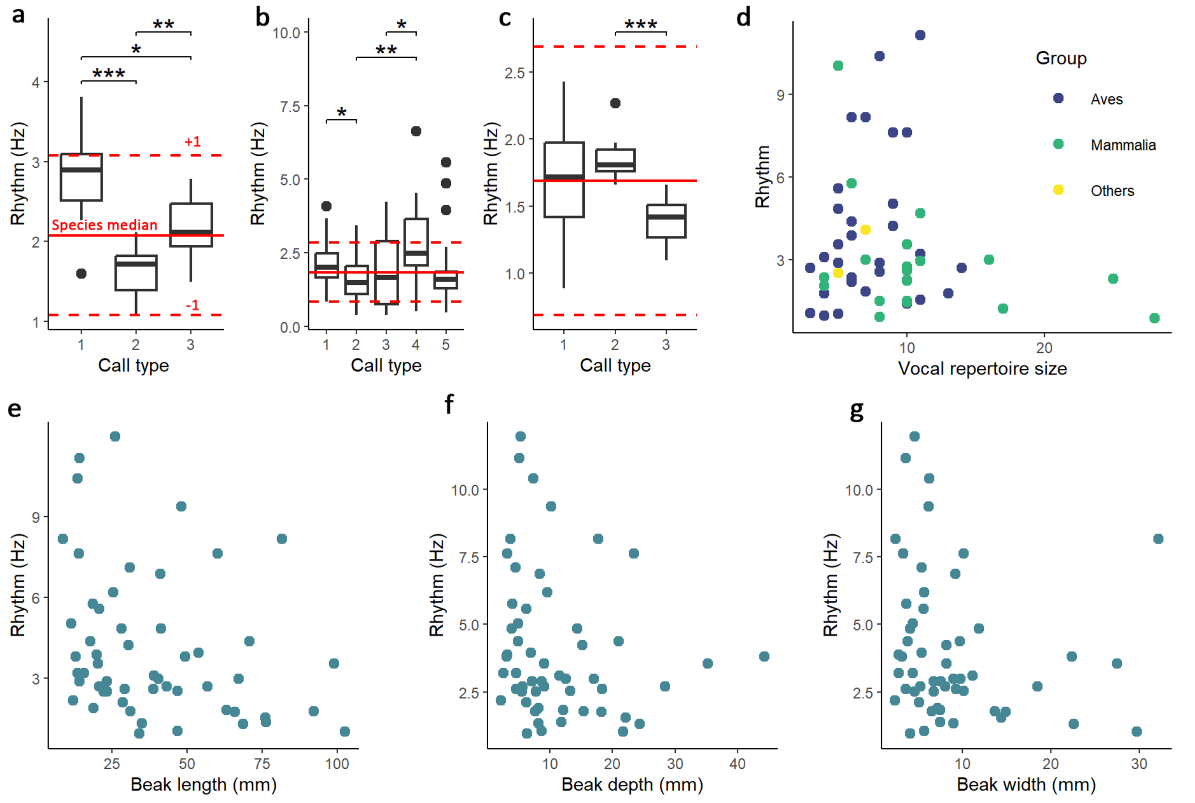


**Supplementary figure 4 Additional Analysis on Rhythm** a) Rhythm in sequences of different call types (1=affiliative grunt, 2=scream, 3=threat grunt) in olive baboons *(Papio anubis)*. b) Rhythm in sequences of different call types (1=bark, 2=growl, 3=howl, 4=snarl,

5= whine) in dogs *(Canis lupus familiaris).* c) Rhythm in sequences of different call types (1=alarm call, 2=flight call, 3=song) in Eurasian stone-curlews *(Burhinus oedicnemus)* d) Rhythm plotted as a function of vocal repertoire size. e) Rhythm plotted as a function of beak length in birds. f) Rhythm plotted as a function of beak depth in birds. g) Rhythm plotted as a function of beak width in birds.
